# Draft genome and transcriptomic sequence data of three invasive insect species

**DOI:** 10.1101/2024.12.02.626401

**Authors:** Eric Lombaert, Christophe Klopp, Aurélie Blin, Gwenolah Annonay, Carole Iampietro, Jérôme Lluch, Marine Sallaberry, Sophie Valière, Riccardo Poloni, Mathieu Joron, Emeline Deleury

**Affiliations:** INRAE, Université Côte d’Azur, ISA, Sophia-Antipolis France; INRAE, Sigenae, MIAT, Castanet Tolosan, France; INRAE, GeT-PlaGe, Genotoul, Castanet-Tolosan, France; CNRS, EPHE, IRD, Université de Montpellier, CEFE, Montpellier, France

**Keywords:** whole-genome sequencing, RNA-seq, insect pest species, *Cydalima perspectalis*, *Leptoglossus occidentalis*, *Tecia solanivora*

## Abstract

*Cydalima perspectalis* (the box tree moth), *Leptoglossus occidentalis* (the western conifer seed bug), and *Tecia solanivora* (the Guatemalan tuber moth) are three economically harmful invasive insect species. This study presents their genomic and transcriptomic sequences, generated through whole-genome sequencing, RNA-seq transcriptomic data, and Hi-C sequencing. The resulting genome assemblies exhibit good quality, providing valuable insights into these species. The genome sizes are 500.4 Mb for *C. perspectalis*, 1.74 Gb for *L. occidentalis*, and 623.3 Mb for *T. solanivora*. These datasets are available in the NCBI Sequence Read Archive (BioProject PRJNA1140410) and serve as essential resources for population genomics studies and the development of effective pest management strategies, addressing significant gaps in the understanding of invasive insect species.

## Background

Despite the recognized threats that invasive species pose to human health, economic activities, and biodiversity, our understanding of the factors driving most biological invasions remains limited (Li et al. 2016; Chinchio et al. 2020; Diagne et al. 2021). This knowledge gap is particularly pronounced for insects, despite them being the most prevalent and destructive group of animal invaders in terrestrial ecosystems (Bradshaw et al. 2016). Many phytophagous insects have a significant impact, attacking crops, ornamental plants, or forest trees of ecological and commercial importance. Yet very few insect genomes are available to address the many fundamental and applied questions related to biological invasions, including insights into genetic adaptations, invasion pathways, and mechanisms underlying their impacts on native ecosystems.

Among the numerous invasive insect species, *Cydalima perspectalis* (the box tree moth), *Leptoglossus occidentalis* (the western conifer seed bug), and *Tecia solanivora* (the Guatemalan tuber moth) are notable for their rapid spread and severe impacts on their respective host plants. *C. perspectalis* is a major defoliator of Buxus species, causing widespread damage to natural and ornamental box plants. Native to East Asia, it was first detected in Europe in 2007 and has since expanded its range to North America (Krüger 2008; Bras et al. 2022; Coyle et al. 2022). *L. occidentalis*, originally from western North America, is a major seed pest of conifers, affecting the regeneration of valuable tree species. It has progressively invaded eastern North America since the 1950s and Europe since 1999, where it has established successful populations (Schaffner 1967; Taylor et al. 2001; Lesieur et al. 2019). *T. solanivora*, a specialist pest of potatoes, has had a devastating impact on potato production in the area it has invaded. Native to Central America, it has spread to South America since the 1970s and was introduced to the Canary Islands in 1999 (Povolny 1973; Puillandre et al. 2008).

Despite their ecological and economic importance, genomic resources for these species remain scarce. Although a genome assembly has been previously published for *C. perspectalis* (Broad et al. 2024), it was generated from an individual of the *typica* morph (light-colored). Here, we provide a new draft genome for this species based on an individual of the *fusca* morph (dark-colored), allowing for comparative genomic studies between these morphs. Furthermore, no reference genome was available for *L. occidentalis* or *T. solanivora*, limiting our ability to study the genetic drivers of their invasions. To fill this gap, we generated draft genome assemblies and transcriptomic data for these three species. These datasets, which cover species with diverse genomic characteristics (Table 1), provide valuable resources for investigating invasion dynamics and developing more effective management strategies.

**Table 1.** Species information and main genome metrics.

| Information and Metrics | *C. perspectalis* | *L. occidentalis* | *T. solanivora* |
| --- | --- | --- | --- |
| Order | Lepidoptera | Hemiptera | Lepidoptera |
| Host plants | *Buxus* species | Conifer species | Potatoes |
| Native area | Eastern Asia | Western North America | Central America |
| Invasive areas | Europe, North America, Middle East | Eastern North America, Europe | South America, Canary Islands |
| Estimated genome size | 468.1 Mb | 1,536.57 Mb | 601.7 Mb |
| Estimated heterozygosity | 1.29% | 1.82% | 1.32% |
| Estimated repeat fraction | 24.1% | 58.3% | 42.1% |
| Assembly accession | GCA_045371335.1 | GCA_045410785.1 | GCA_045412185.1 |

## Data description and validation

### Raw read quality

Long-reads, transcriptome RNA-seq and Hi-C sequences were obtained via PacBio and Illumina platforms. Raw reads were deposited onto the NCBI Sequence Read Archive under BioProject PRJNA1140410. All BioSample and SRA accession numbers are listed in Table 2. Large quantities of reads were produced, and the sequence qualities were high for all libraries (Table 2). Mapping the reads from each library onto the genome assemblies built from these same data resulted in good alignment rates for long reads (nearly 100%) and Hi-C sequences (close to 97%). For RNA-seq, mapping rates were more variable, with an average of 72.5% and some individuals exhibiting low values (see Table 2 for details).

**Table 2.** Read set statistics, including quality evaluation. Cp = *Cydalima perspectalis*; Lo = *Leptoglossus occidentalis*; Ts = *Tecia solanivora*.

| BioSample accession number | SRA accession number | Species | Strategy | Read count | Nuc count | Avg qual | Alignment rate (%) |
| --- | --- | --- | --- | --- | --- | --- | --- |
| SAMN42831079 | SRR30002755 | Cp | WGS | 2,336,698 | 35,334,168,532 | 82.27 | 99.97 |
| SAMN42831080 | SRR30002754 | Cp | RNA-seq | 74,625,830 | 11,105,068,576 | 35.60 | 82.67 |
| SAMN42831081 | SRR30002753 | Cp | RNA-seq | 58,079,426 | 8,628,250,664 | 35.65 | 74.15 |
| SAMN42831082 | SRR30002764 | Cp | RNA-seq | 71,460,408 | 10,610,626,515 | 35.60 | 68.61 |
| SAMN42831083 | SRR30002763 | Cp | RNA-seq | 54,014,740 | 8,019,510,478 | 35.60 | 84.22 |
| SAMN42831084 | SRR30002762 | Cp | RNA-seq | 125,597,334 | 18,795,890,549 | 35.55 | 85.02 |
| SAMN42831085 | SRR30002761 | Cp | RNA-seq | 66,358 | 9,926,612 | 35.65 | 84.51 |
| SAMN42831072 | SRR30002766 | Lo | WGS | 1,951,868 | 36,504,448,466 | 79.14 | 99.96 |
| SAMN42831073 | SRR30002765 | Lo | RNA-seq | 44,803,922 | 6,693,029,521 | 35.55 | 58.32 |
| SAMN42831074 | SRR30002760 | Lo | RNA-seq | 77,860,552 | 11,532,876,985 | 35.70 | 77.17 |
| SAMN42831075 | SRR30002759 | Lo | Hi-C | 90,074,518 | 13,511,177,700 | 35.56 | 96.98 |
| SAMN42831076 | SRR30002758 | Ts | WGS | 1,260,730 | 14,430,962,828 | 84.60 | 99.98 |
| SAMN42831077 | SRR30002757 | Ts | RNA-seq | 59,597,426 | 8,895,621,390 | 35.55 | 36.92 |
| SAMN42831078 | SRR30002756 | Ts | RNA-seq | 94,062,578 | 13,956,137,243 | 35.65 | 73.33 |

### Genome assemblies and quality control

For each of the three species, *C. perspectalis, L. occidentalis* and *T. solanivora*, draft de novo genomes were assembled. Hi-C sequencing enhanced scaffolding for *C. perspectalis* (public read set: ERR11217097) and *L. occidentalis* (read set from this study). N50 values indicate a high level of contiguity for all three assemblies, exceeding 15 Mb in each case. *C. perspectalis* had the least fragmented assembly, with a total length of 469.1 Mb, only 52 scaffolds and a high sequencing depth of 75X (Table 3). Despite its larger genome size (1.77 Gb) and lower sequencing depth (22.5X), *L. occidentalis* exhibits strong contiguity, as reflected by its high N50 value of 147.7 Mb (Table 3). Additional quality indicators, including BUSCO scores and Mercury QV metrics, further validated the overall high quality of all three assemblies (Table 3).

**Table 3.**
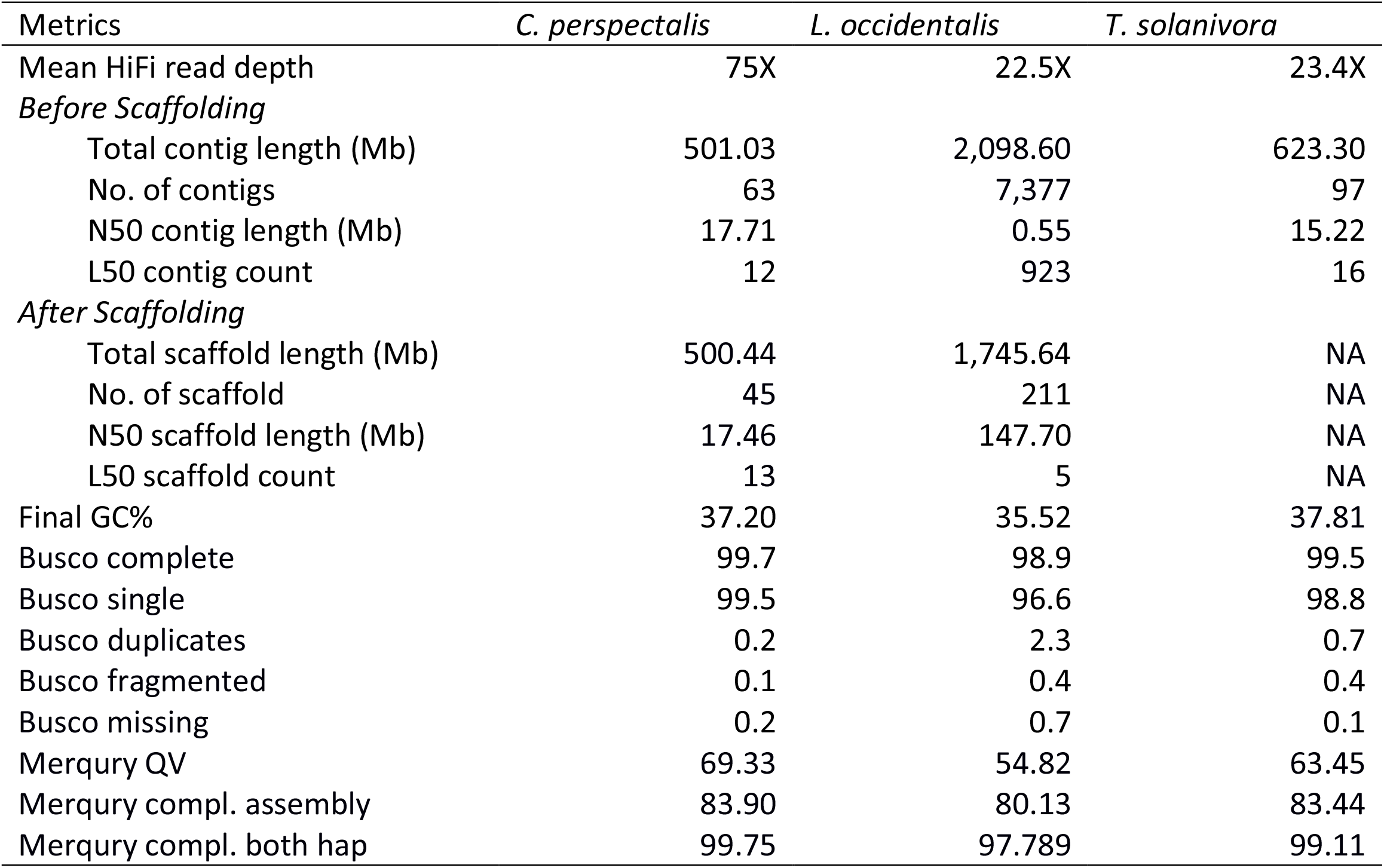
Genome assembly metrics, including quality control metrics.

| Metrics | *C. perspectalis* | *L. occidentalis* | *T. solanivora* |
| --- | --- | --- | --- |
| Mean HiFi read depth | 75X | 22.5X | 23.4X |
| *Before Scaffolding* |  |  |  |
| Total contig length (Mb) | 501.03 | 2,098.60 | 623.30 |
| No. of contigs | 63 | 7,377 | 97 |
| N50 contig length (Mb) | 17.71 | 0.55 | 15.22 |
| L50 contig count | 12 | 923 | 16 |
| *After Scaffolding* |  |  |  |
| Total scaffold length (Mb) | 500.44 | 1,745.64 | NA |
| No. of scaffold | 45 | 211 | NA |
| N50 scaffold length (Mb) | 17.46 | 147.70 | NA |
| L50 scaffold count | 13 | 5 | NA |
| Final GC% | 37.20 | 35.52 | 37.81 |
| Busco complete | 99.7 | 98.9 | 99.5 |
| Busco single | 99.5 | 96.6 | 98.8 |
| Busco duplicates | 0.2 | 2.3 | 0.7 |
| Busco fragmented | 0.1 | 0.4 | 0.4 |
| Busco missing | 0.2 | 0.7 | 0.1 |
| Merqury QV | 69.33 | 54.82 | 63.45 |
| Merqury compl. assembly | 83.90 | 80.13 | 83.44 |
| Merqury compl. both hap | 99.75 | 97.789 | 99.11 |

Consistent with the relatively high heterozygosity observed in all three genomes (Table 1), K-mer spectra-cn plots reveal two distinct peaks based on HiFi reads, each represented once in the assembly (Fig. 1). In the genomes of *L. occidentalis* and *T. solanivora*, slight additional k-mer fractions appear more than once in the assembly (Fig. 1b and Fig. 1c), suggesting incomplete removal of the second haplotype. However, due to the low sequencing depth for both species, the possibility of ancient duplications cannot be ruled out.

**Figure 1.**
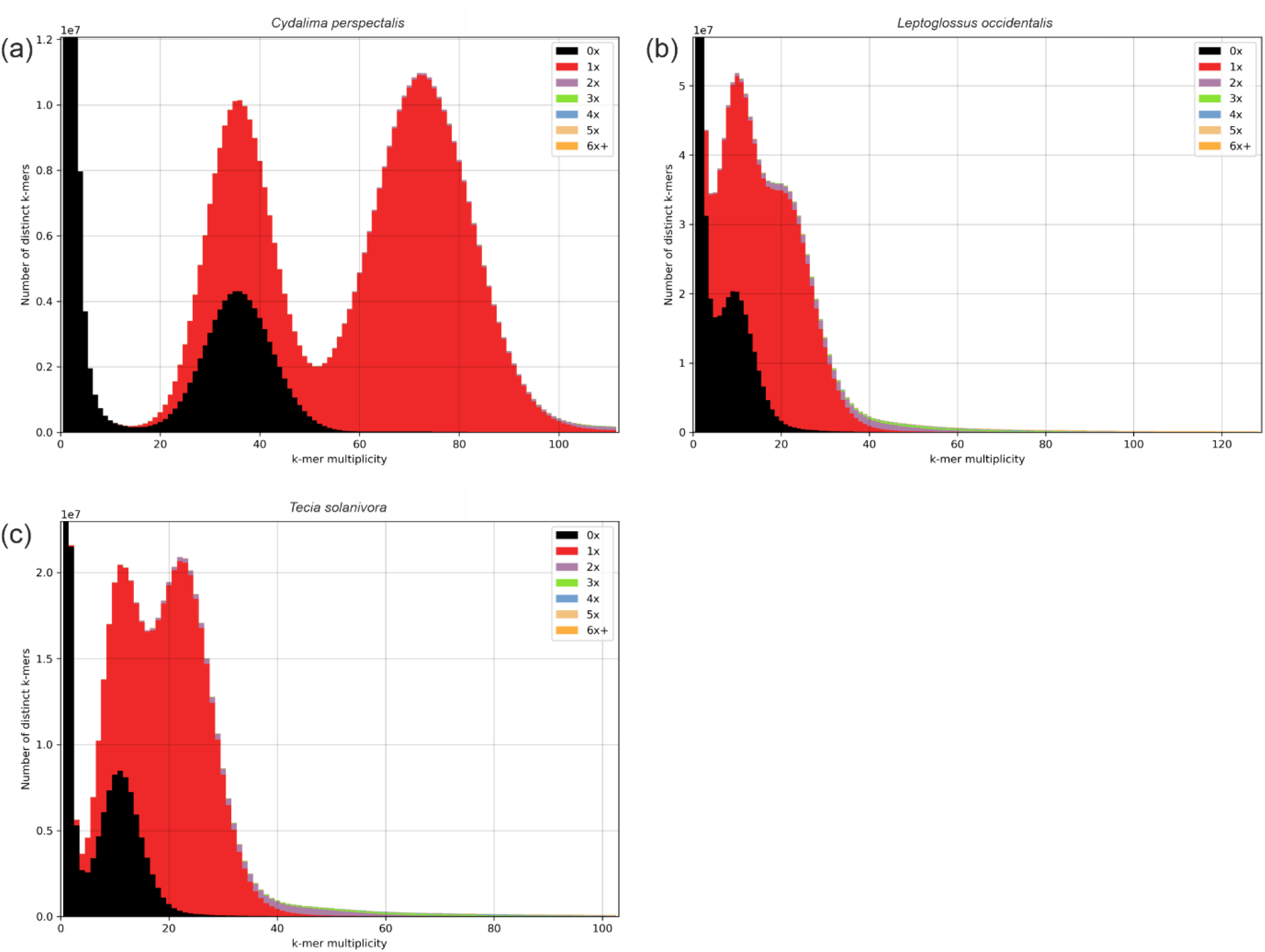
HiFi reads k-mer spectra-cn plots for (a) *Cydalima perspectalis*, (b) *Leptoglossus occidentalis*, and (c) *Tecia solanivora*. The x-axis represents k-mer multiplicity, while the y-axis indicates the count of distinct k-mers at a given coverage. The presence of two main peaks in all three species reflects heterozygosity, with the first peak corresponding to heterozygous k-mers and the second to homozygous k-mers.

Alignments against closely related reference genomes yielded mixed results (Fig. 2). For *C. perspectalis*, the alignments showed high concordance between scaffolds and the reference chromosome of the same species across the entire genome (Fig. 2a). By contrast, *L. occidentalis* and *T. solanivora* were aligned with more distantly related species, *Leptoglossus phyllopus* and *Teleiodes luculella*, respectively, resulting in lower overall concordance as expected (Fig. 2b and Fig. 2c). Nonetheless, large blocks of collinearity were observed for both species, confirming the strong accuracy and contiguity of our assemblies.

**Figure 2.**
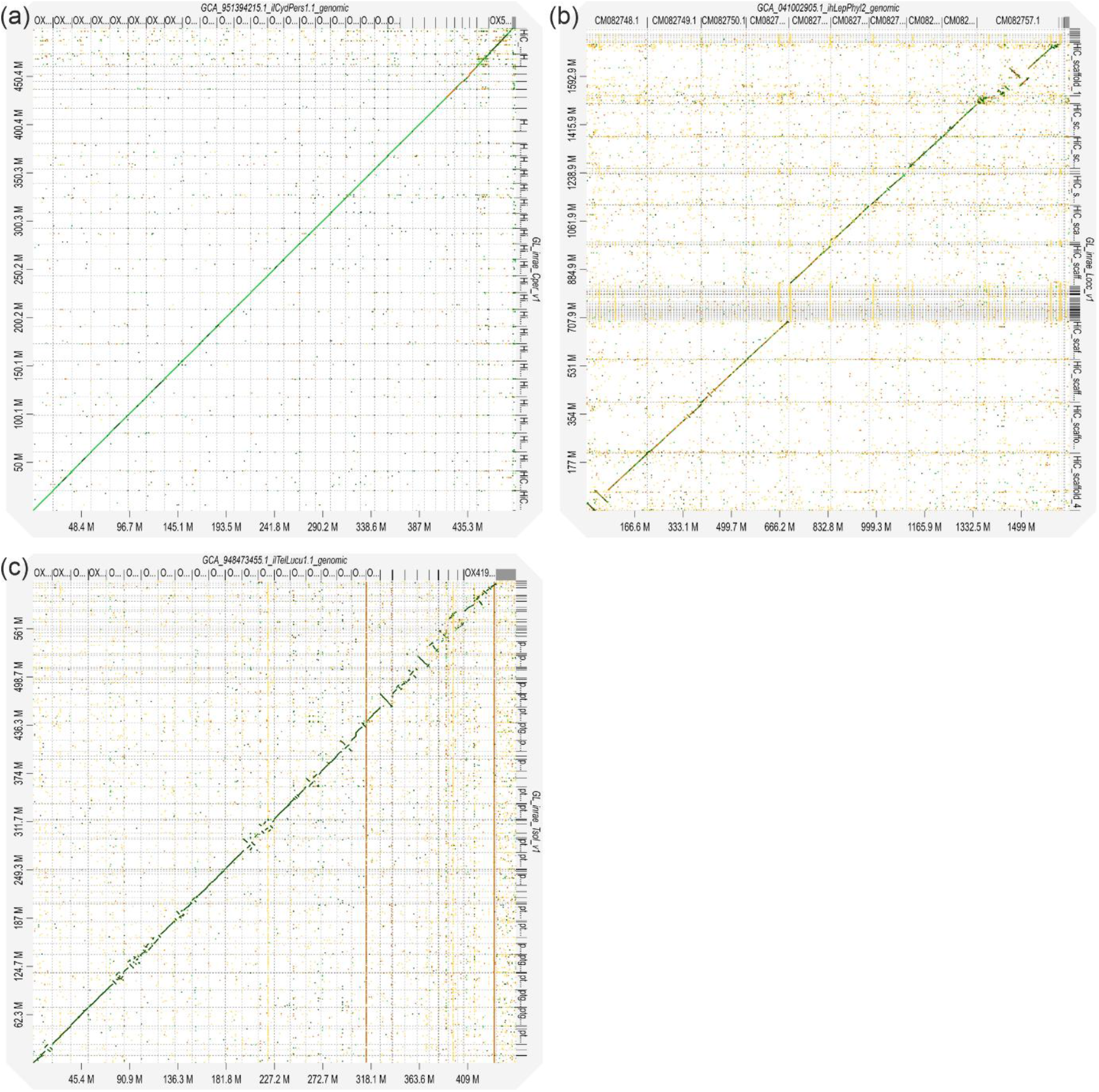
Dot-plots showing whole-genome alignment with minimap2 of each of the three assembled genomes (Y-axis) to publicly available genomes of related species (X-axis): (a) the assembly resulting from this study of *Cydalima perspectalis* versus the reference genome of the same species (GenBank reference: GCA_951394215.1); (b) the assembly of *Leptoglossus occidentalis* versus the reference genome of *Leptoglossus phyllopus* (GenBank reference: GCA_041002905.1); (c) the assembly of *Tecia solanivora* versus the reference genome of *Teleiodes luculella* (GenBank reference: GCA_948473455.1).

Overall, our assemblies show high contiguity and completeness, despite the presence of heterozygosity and repeat content (Table 1). The use of HiFi long reads, combined with Hi-C scaffolding where available, allowed us to mitigate these challenges and produce high-quality genomic resources.

### Genome annotation and annotation quality control

Annotation across all three genomes predicted a similar number of genes, ranging from 19,326 to 20,895 (Table 4). BUSCO scores were consistently high, with the lowest completeness score reaching 96.0% (Table 4).

**Table 4.** Genome annotation quality control metrics.

| Metrics | *C. perspectalis* | *L. occidentalis* | *T. solanivora* |
| --- | --- | --- | --- |
| Annotation procedure | Helixer + Braker + agat | Helixer | Helixer + Braker + agat |
| Number of genes | 19,326 | 20,895 | 20,019 |
| Number of transcripts | 19,370 | 20,895 | 24,957 |
| Number of exons | 132,873 | 150,750 | 297,593 |
| Busco complete | 98.5 | 96.0 | 96.5 |
| Busco single | 98.3 | 94.6 | 96.1 |
| Busco duplicates | 0.2 | 1.4 | 0.4 |
| Busco fragmented | 1.0 | 1.7 | 1.5 |
| Busco missing | 0.5 | 2.3 | 2.0 |

## Methods

### Sample collection and extraction

All individuals were collected and extracted between March and September 2022 (Table 5). *C. perspectalis* samples were obtained from a rearing facility (in CEFE, Montpellier, France), *L. occidentalis* samples were collected from pine trees in southern France, and *T. solanivora* samples were collected from a potato field in central Colombia. Before further processing, the *C. perspectalis* sample used for long-read sequencing was stored dry at −80°C, while those used for RNA-seq were stored in RNALater at −20°C. *L. occidentalis* and *T. solanivora* samples were stored dry at −80°C and in RNALater at −20°C, respectively.

**Table 5.**
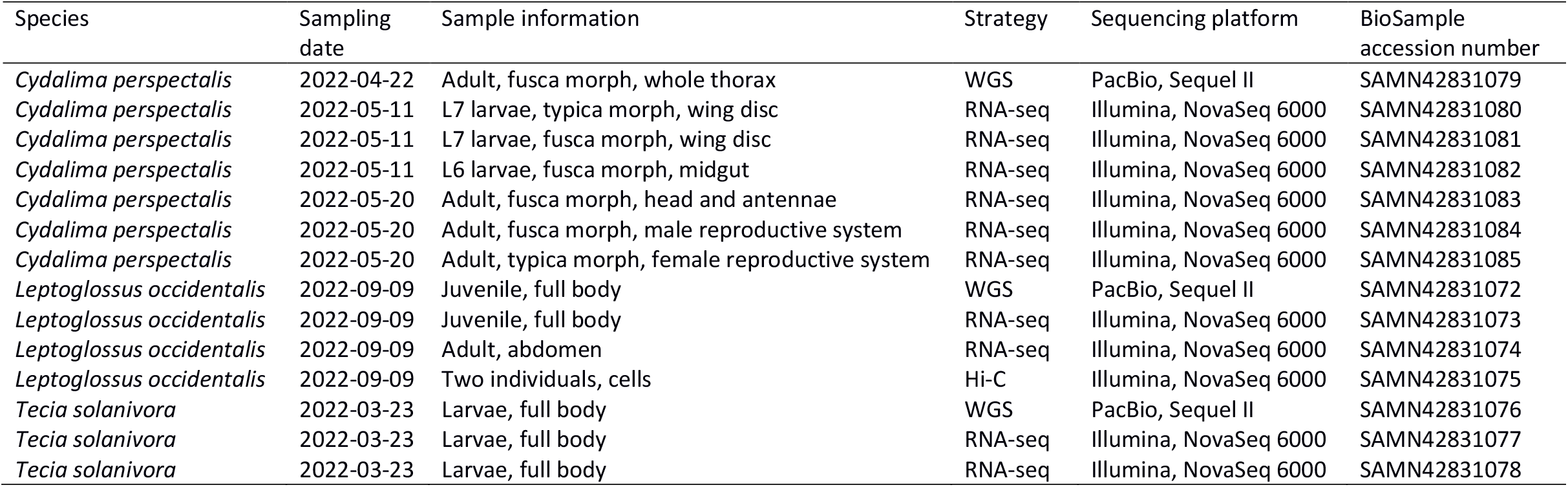
Sample information and sequencing methods.

| Species | Sampling date | Sample information | Strategy | Sequencing platform | BioSample accession number |
| --- | --- | --- | --- | --- | --- |
| *Cydalima perspectalis* | 2022-04-22 | Adult, fusca morph, whole thorax | WGS | PacBio, Sequel II | SAMN42831079 |
| *Cydalima perspectalis* | 2022-05-11 | L7 larvae, typica morph, wing disc | RNA-seq | Illumina, NovaSeq 6000 | SAMN42831080 |
| *Cydalima perspectalis* | 2022-05-11 | L7 larvae, fusca morph, wing disc | RNA-seq | Illumina, NovaSeq 6000 | SAMN42831081 |
| *Cydalima perspectalis* | 2022-05-11 | L6 larvae, fusca morph, midgut | RNA-seq | Illumina, NovaSeq 6000 | SAMN42831082 |
| *Cydalima perspectalis* | 2022-05-20 | Adult, fusca morph, head and antennae | RNA-seq | Illumina, NovaSeq 6000 | SAMN42831083 |
| *Cydalima perspectalis* | 2022-05-20 | Adult, fusca morph, male reproductive system | RNA-seq | Illumina, NovaSeq 6000 | SAMN42831084 |
| *Cydalima perspectalis* | 2022-05-20 | Adult, typica morph, female reproductive system | RNA-seq | Illumina, NovaSeq 6000 | SAMN42831085 |
| *Leptoglossus occidentalis* | 2022-09-09 | Juvenile, full body | WGS | PacBio, Sequel II | SAMN42831072 |
| *Leptoglossus occidentalis* | 2022-09-09 | Juvenile, full body | RNA-seq | Illumina, NovaSeq 6000 | SAMN42831073 |
| *Leptoglossus occidentalis* | 2022-09-09 | Adult, abdomen | RNA-seq | Illumina, NovaSeq 6000 | SAMN42831074 |
| *Leptoglossus occidentalis* | 2022-09-09 | Two individuals, cells | Hi-C | Illumina, NovaSeq 6000 | SAMN42831075 |
| *Tecia solanivora* | 2022-03-23 | Larvae, full body | WGS | PacBio, Sequel II | SAMN42831076 |
| *Tecia solanivora* | 2022-03-23 | Larvae, full body | RNA-seq | Illumina, NovaSeq 6000 | SAMN42831077 |
| *Tecia solanivora* | 2022-03-23 | Larvae, full body | RNA-seq | Illumina, NovaSeq 6000 | SAMN42831078 |

### Long-read DNA sequencing

High-molecular-weight DNA was extracted from one individual of each species. The QIAGEN Genomic Tip 100/G kit was used for *C. perspectalis*, and the PROMEGA Wizard Genomic DNA purification kit for *L. occidentalis* and *T. solanivora*.

Library preparation and sequencing were performed at GeT-PlaGe core facility, INRAe Toulouse according to the manufacturer’s instructions “Procedure & Checklist – Preparing whole genome and metagenome libraries using SMRTbell® prep kit 3.0”. At each step, DNA was quantified using the Qubit dsDNA HS Assay Kit (Life Technologies). DNA purity was tested using the nanodrop (Thermofisher) and size distribution and degradation assessed using the Femto pulse Genomic DNA 165 kb Kit (Agilent). Purification steps were performed using AMPure PB beads (PacBio) and SMRTbell cleanup beads (Pacbio). We made a DNA repair step with the “SMRTbell Damage Repair Kit SPV3” (PacBio). DNA was purified then sheared using the Megaruptor1 system (Diagenode) (Only the Te91 sample was not sheared). After a nuclease step using “SMRTbell® prep kit 3.0”, the libraries were size-selected, using a cutoff on the Pippin HT Size Selection system (Sage Science) with “0.75% Agarose, 6-10 kb High Pass, 75E” protocol.

Using Binding kit 3.2 (primer 3.2, polymerase 2.2) and sequencing kit 2.0, the libraries were sequenced by Adaptive Loading onto three SMRTcells (one per species) on Sequel2 instrument at 90 pM with a 2-hour pre-extension and a 30-hour movie.

### *Hi-C sequencing of* L. occidentalis

For *L. occidentalis*, Hi-C was performed using the Arima High Coverage Hi-C kit (reference number A101030) according to the manufacturer’s instructions. Briefly, tissues from two frozen adult individuals were crosslinked and homogenized. Chromatin was then digested using four restriction enzymes. Chromatin ends were biotinylated and ligated by proximity ligation. After reverse crosslink and purification, the DNA was used for library preparation using Arima Library Prep Module. Sequencing library was performed using the Arima Library Prep (reference number A303010). The Hi-C library was sequenced on an Illumina NovaSeq6000 platform to generate 2 x150 bp read pairs.

### Transcriptomic data sequencing

We used the QIAGEN RNeAsy mini plus kit to extract RNA from various dissected tissues of three larvae and three adults of *C. perspectalis*, full bodies of one juvenile and one adult of *L. occidentalis*, and full bodies of two larvae of *T. solanivora* (see Table 5 for details).

RNAseq was performed at the GeT-PlaGe core facility, INRAe Toulouse. RNA-seq libraries were prepared according to Illumina’s protocols using the Illumina TruSeq Stranded mRNA sample prep kit to analyze mRNA. Briefly, mRNA was selected using poly-T beads. RNA was then fragmented to generate double stranded cDNA and adaptors were ligated to be sequenced. 11 cycles of PCR were applied to amplify libraries. Library quality was assessed using a Fragment Analyser and libraries were quantified by QPCR using the Kapa Library Quantification Kit. RNA-seq experiments have been performed on an Illumina NovaSeq 6000 using a paired-end read length of 2×150 pb with the Illumina NovaSeq 6000 sequencing kits.

### Sequence quality validation

Read quality was checked with two methods. First by computing the average base pair quality value. Second by aligning the reads to the assembly and checking the rate of primary mappings. RNA-seq reads were preprocessed using fastp version 0.23.4 (Chen et al. 2018) with default parameters before alignment. Alignments were performed with minimap2 version 2.24 (Li 2018) for long reads (with the -x map-hifi parameter) and Hi-C reads (−x sr parameter), and with STAR v2.7.11b (Dobin et al. 2013) for RNA-seq reads. The alignment files were transformed into BAM format with samtools sort version 1.14 (Li et al. 2009), and the alignment metrics were extracted using samtools flagstat with default parameters.

### Genome assemblies and scaffolding

All three assemblies were generated using hifiasm (Cheng *et al*., 2021, 2022). Due to the differing data availability timelines, *Cydalima perspectalis* reads were assembled with version 0.16.1, while *Leptoglossus occidentalis* and *Tecia solanivora* were assembled with version 0.18.8, applying default parameters in all cases. Before scaffolding, the assembly of *Leptoglossus occidentalis* was processed with purge_dups version 1964aaa using default parameters to remove large k-mer duplications. This reduced the total assembly size from 2.098 Gb to 1.769 Gb and the number of contigs from 7,377 to 4,976.

*C. perspectalis* and *L. occidentalis* were scaffolded into chromosomes using public and novel Hi-C reads respectively. The public *C. perspectalis* hi-C read set ERR11217097 was downloaded from the NCBI. Both scaffoldings were performed using juicer version 1.6, 3D-DNA release 529ccf4 (Dudchenko et al. 2017), followed by a manual curation with Juicebox version 1.11.08 (Durand et al. 2016).

Additionally, all three mitochondrial genomes were assembled using MitoHiFi v2.2 (Uliano-Silva et al. 2023).

### Genome assemblies validation

The assemblies in their final states were validated with four methods. First assembly metrics were calculated using assemblathon_stats.pl script (Bradnam et al. 2013). Second BUSCO scores were calculated using the insecta_odb10 database (75 genomes and 1367 BUSCOs) with version 5.7.1 of the BUSCO software package (Manni et al. 2021), with the -m geno option. Third merqury QV and completeness statistics were generated (Rhie et al. 2020). Fourth, d-genies dot-plots (Cabanettes and Klopp 2018) were generated versus a phylogenetically close related species (itself in the case of *Cydalima perspectalis* for which a reference genome is available).

### Genome metrics

Genome metrics corresponding to estimated genome size, repeat fraction and heterozygosity found in Table 1 were all extracted from the HiFi reads using genomescope2 (Ranallo-Benavidez et al. 2020). The *Cydalima perspectalis* estimated genome size is very close to the size of the good quality public assembly found at the NCBI GCA_951394215.1 (483.7 Mb).

### Genome annotation and annotation validation

Using the available RNA-Seq reads sets, we performed a de novo annotation for each assembly, primarily to validate nucleotide content of the assemblies. Annotations were conducted with helixer version 0.3.1_cuda_11.2.0 (Stiehler et al. 2020) using the invertebrate trained model and default parameters, as well as with braker (Gabriel et al. 2021). Both annotations were then merged using the agat_sp_merge_annotations.pl script from the agat suite version 1.2.0 with default parameters (Dainat et al. 2021). Annotation validation was performed using the same version of BUSCO software as for the assemblies but with the -m tran option. Due to poor results from braker for *L. occidentalis*, only helixer output was kept as final annotation for this species.

Additionally, all three mitochondrial genomes were annotated using GeSeq v2.03 (Tillich et al. 2017).

## Acknowledgments

We thank our colleagues Stéphane Dupas and Barbara Porro for *Tecia solanivora* and *Leptoglossus occidentalis* samples. We thank the Technical Platform for Experimental Ecology at CEFE Montpellier for their assistance in maintaining captive *C. perspectalis* stocks. We also thank Emmanuelle Murciano-Germain for administrative assistance. This work was performed in collaboration with the GeT core facility, Toulouse, France (GeT, https://doi.org/10.15454/1.5572370921303193E12). GeT core facility was supported by France Génomique National infrastructure, funded as part of “Investissement d’avenir” program managed by Agence Nationale pour la Recherche (contract ANR-10-INBS-09).

## Data availability

The sequencing data and genome assemblies from this study have been deposited at DDBJ/ENA/GenBank under project PRJNA1140410: https://www.ncbi.nlm.nih.gov/bioproject/1140410

Genome and mitogenome annotations, as well as mitogenome sequences, are available at Data INRAE via the following link: https://doi.org/10.57745/WDMFPB

All the required command-line instructions can be accessed via the following link: https://github.com/chklopp/FARDOMICS_assemblies/blob/main/README.md

## Author contributions

EL, CK and ED designed the study. EL, RP and MJ managed the choice and collection of samples. AB and RP prepared the samples. GA, CI, JL, MS and SV prepared the libraries and performed the sequencing. CK and EL analysed the data. EL, CK, CI, MS and SV wrote the paper. All authors have revised and approved the final manuscript.

